# Acoelomorph flatworm monophyly is a severe long branch-attraction artefact obscuring a clade of Acoela and Xenoturbellida

**DOI:** 10.1101/2023.08.27.555014

**Authors:** Anthony K Redmond

## Abstract

Acoelomorpha is a broadly accepted clade of bilaterian animals made up of the fast-evolving, morphologically simple, mainly marine flatworm lineages Acoela and Nemertodermatida. Phylogenomic studies support Acoelomorpha’s close relationship with the slowly evolving and similarly simplistic *Xenoturbella*, together forming the phylum Xenacoelomorpha. The phylogenetic placement of Xenacoelomorpha amongst bilaterians is controversial, with some studies supporting Xenacoelomorpha as the sister group to all other bilaterians, implying that their simplicity may be representative of early bilaterians. Others propose that this placement is a long branch attraction artefact resulting from the fast-evolving Acoelomorpha, and instead suggest that they are the secondarily simplified sister group of the deuterostome clade Ambulacraria. Perhaps as a result of this debate, internal xenacoelomorph relationships have been somewhat overlooked at a phylogenomic scale. Here, I employ both empirical and simulation approaches to detect and overcome phylogenomic errors to reassess the relationship between *Xenoturbella* and the fast evolving acoelomorph flatworms. I conclude that subphylum Acoelomorpha is a long-branch attraction artefact obscuring a previously undiscovered clade comprising *Xenoturbella* and Acoela, for which I propose the name Xenacoela. These analyses are also consistent with the Nephrozoa hypothesis deriving from systematic error, and instead generally favour a close, but unclear, relationship of Xenacoelomorpha with deuterostomes. This study provides a template for future efforts aimed at discovering and correcting unrecognised long-branch attraction artefacts throughout the tree of life.

## Introduction

Xenacoelomorpha is an enigmatic, typically marine, phylum of invertebrate bilaterian animals^1–4^. They are characterised by apparently simple morphology, particularly their acoelomate body plan and the absence of nephridia, but also lack characteristic features found in many bilaterians such as a through-gut, circulatory and respiratory systems, and a complex brain^1,2,4^. However, recent studies focused on the lineage have revealed remarkable diversity in nervous system morphology^5–7^, as well as evidence for active excretion despite the lack of a specialised organ^8,9^, implying underappreciated biological complexity in these species. The lineage is divided into two subphyla^10^; the fast evolving Acoelomorpha^11^, consisting of the two acoelomorph flatworm classes Acoela and Nemertodermatida, and the more slowly evolving Xenoturbellida^3,12^, from which only the genus *Xenoturbella* is known.

Although morphological similarity has long been noted between *Xenoturbella* and Acoelomorpha^13–16^, their confident joining within Xenacoelomorpha is a relatively recent phylogenomic discovery^3,17–19^. Prior to this, early studies considered both as “turbellarian” flatworms, but both lineages proved generally difficult to place amongst Bilateria^4,11,15,17,20,20–24^. Nonetheless, while phylogenomics strongly supports Xenacoelomorpha, and dismisses a close relationship with platyhelminths (including other “Turbellaria”)^3,17,18,22^, consensus on the relationship of Xenacoelomorpha to other Bilateria has not been reached. Some studies favour Xenacoelomorpha as the sister group to all other Bilaterians (the Nephrozoa hypothesis)^17,18,25^, suggesting that their relatively simple morphology is representative of early bilaterians (**Fig. 1A**). Others argue that Xenacoelomorpha is the sister group to Ambulacraria (the Xenambulacraria hypothesis)^3,26–28^, implying that their morphology is degenerate (**Fig. 1A**). In the latter case Nephrozoa is viewed to be the result of systematic error stemming from the fast-evolving and long-branching Acoelomorpha^3,26–30^. Additionally, some analyses have recovered other placements, such as sister to either deuterostomes^3^ or protostomes^25^. Perhaps most remarkably, recent efforts to resolve this problem by minimising systematic error have questioned the monophyly of deuterostomes^26,31^.

**Figure 1.**
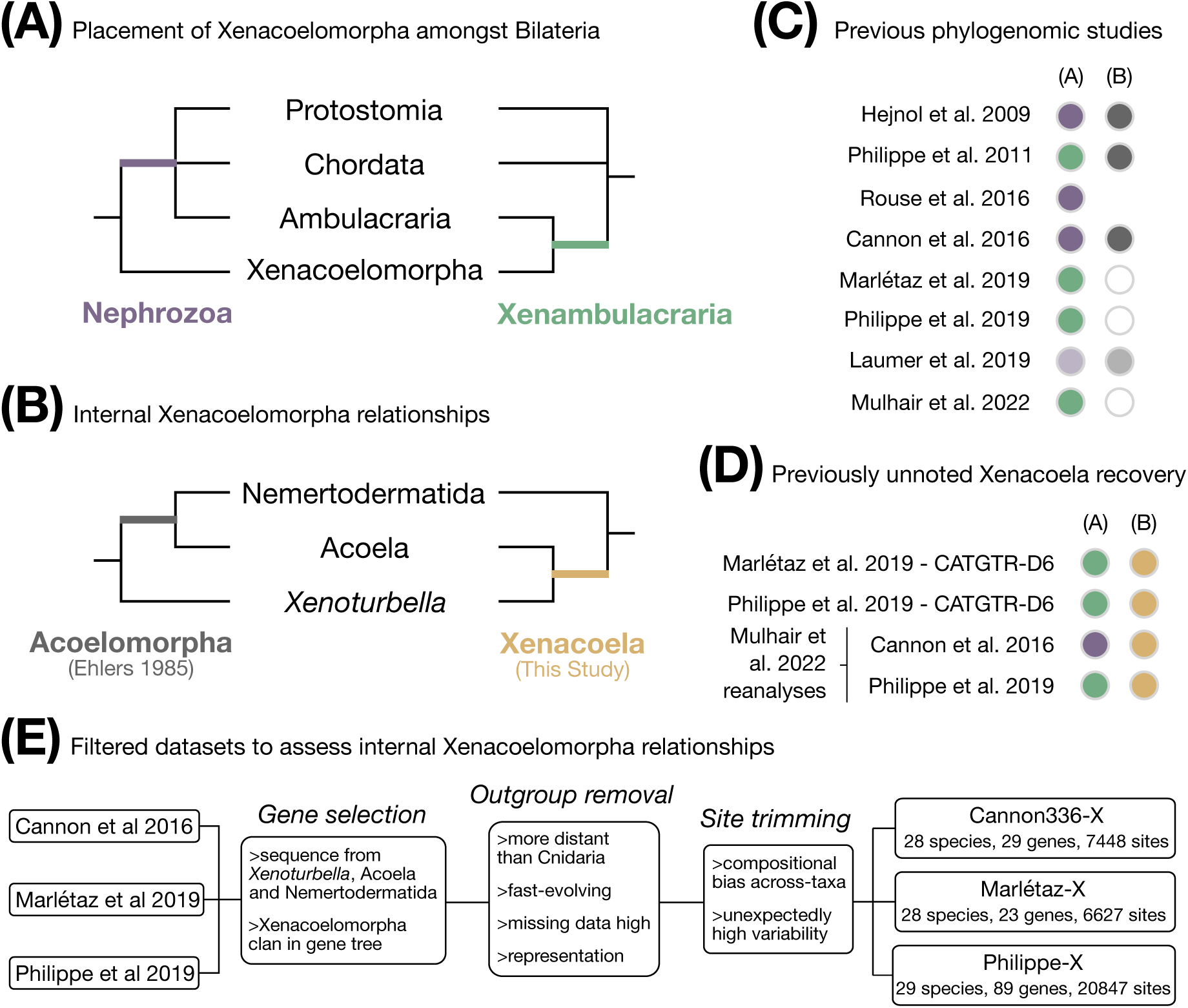
Hypothesized Xenacoelomorpha relationships and dataset preparation. (**A**) Conflicting hypotheses for the placement of Xenacoelomorpha within Bilaterian evolution as either the sister group to all other bilaterians (Nephrozoa; branch shown in purple) or as sister to Ambulacraria (Xenambulacraria; branch shown in green). Relationships between Chordata, Protostomia and (Xen)Ambulacraria are shown as a polytomy as the monophyly of Deuterostomia (Chordata+[Xen]Ambulacraria) has been questioned by recent studies^26,31^. **(B)** Relationships within Xenacoelomorpha. Acoelomorpha (branch shown in grey), the generally accepted hypothesis is shown on the left, while the proposed alternative Xenacoela (branch shown in gold) is shown to the right. **(C)** Conclusions from key past phylogenomic studies are shown with the specific relationship in question and colours are referring to and parts **(A)** and **(B)**. White filled circles indicate that internal Xenacoelomorpha relationships were not discussed. For Laumer et al. 2019, where the Nephrozoa and Acoelomorpha topologies were recovered in all analyses, lower colour saturation for these topologies is used as Xenacoelomorpha was ‘not addressed’ in the study. **(D)** Unreported recovery of Xenacoela in key analyses in previous studies (CATGTR: model used, D6: data were Dayhoff6 recoded; Mulhair et al. reanalyses: data analysed under CATGTR using only a subset of genes for which there is good recovery of indisputable animal clades in gene trees). The specific relationship in question and colour for supported topology are referring to are from parts **(A)** and **(B)**. Note that this is only a subset of the analyses performed in these studies. **(E)** Simple breakdown of the data filtering approach used to produce datasets targeted at internal Xenacoelomorpha relations and with reduced propensity for systematic error.

This ongoing problem highlights the importance of the underlying methodology (e.g., orthology assignment, modelling strategy, etc.) applied in phylogenomics. Past studies have shown that compositional heterogeneity across sites and taxa are important biasing factors in resolving bilaterian relationships using phylogenomic approaches^26–28^. Strategies to reduce such heterogeneity include the use of the CATGTR model^32^, which is designed to accommodate heterogeneity across sites^32,33^, and/or recoding (i.e. binning) of amino acids into a smaller alphabet based on their evolutionary or biochemical properties (e.g. the 6 Dayhoff categories^34^), which can reduce heterogeneity across both sites and taxa but may also risk masking informative substitutions^30,35–40^. Hidden paralogy, where non-orthologous genes are unintentionally incorporated into phylogenomic datasets, as well as other data errors, can also mislead phylogenomic analyses^28,41,42^. Recent studies suggest that such data and modelling errors may bias phylogenomic results towards Nephrozoa, and that when efforts are made to carefully select genes with strong orthologous signals and limit the impact of compositional heterogeneity support for Xenambulacraria emerges^26–29^.

Given the importance to understanding bilaterian evolution, resolving their placement among other animals has understandably been the main goal of phylogenomic analyses including Xenacoelomorpha^18,25,26,28^. Combined with such studies generally and expectedly recovering monophyletic Acoela, Nemertodermatida, Acoelomorpha, and *Xenoturbella*^3,18,26^, and the paucity of genome data for the lineage (although this is beginning to change^9,26,43–45^), this has resulted in little phylogenomic focus on internal xenacoelomorph relationships. Here, using both empirical and simulation approaches intended to detect and overcome phylogenomic error, I reassess the relationships between *Xenoturbella* and the fast evolving acoelomorph flatworms. I conclude that Acoelomorpha is a long-branch attraction artefact obscuring a clade comprising *Xenoturbella* and Acoela, which I tentatively name ‘Xenacoela’ (**Fig. 1B**). Furthermore, the results suggest the Nephrozoa hypothesis of bilaterian evolution is also a phylogenetic error.

## Results

### Unacknowledged support for Xenacoela in past studies

Since the formal proposal of Xenacoelomorpha by Philippe et al (2011)^3^, only two studies focused on the relationships of Xenacoelomorpha have generated large-scale phylogenomic datasets and included members of the three major lineages. These are Cannon et al (2016)^18^, which was built from transcriptomic data and supported Nephrozoa, and Philippe et al (2019)^26^, based mainly on genomic data and supported Xenambulacraria. Other recent studies either did not include Nemertodermatida (e.g. Rouse et al. (2016)^25^), or have employed reanalyses of existing datasets (Mulhair et al. 2022)^28^. However, studies targeting other relationships have occasionally included the three major xenacoelomorph lineages, including that of Marlétaz et al (2019)^27^, which notably includes two *Xenoturbella* species, and that of Laumer and colleagues (2019)^46^.

As a first step to reassessing the relationships between *Xenoturbella* and the acoelomorph flatworms I performed a closer inspection of the phylogenies produced in the aforementioned studies. While most past analyses support the expected sister group relationship between *Xenoturbella* and Acoelomorpha, including all analyses in Cannon et al (2016)^18^, it is notable that the more recent studies did not address internal Xenacoelomorpha relationships^26–28^ (**Fig. 1C**). This may in part be because the Acoelomorpha grouping was not considered to be in doubt, yet key analyses in these studies have sometimes recovered the newly proposed clade Xenacoela (**Fig. 1D**). Importantly, this clade tends to be recovered when efforts to minimise phylogenetic error are applied, e.g., using site-heterogeneous models (such as CATGTR), amino acid recoding, or filtering genes with poor orthologous signal. Specifically, CATGTR analyses in Philippe et al (2019)^26^ did not recover strong support for Acoelomorpha, while combining CATGTR with recoding recovered strong support for Xenacoela (**Fig. 1D**). Philippe et al (2019)^26^ also reanalysed the main Cannon et al (2016)^18^ dataset using CATGTR with recoding and this failed to recover strong support for Acoelomorpha (**Fig. 1D**). Marlétaz et al (2019)^27^ also recovered Xenacoela when combining CATGTR with recoding (**Fig. 1D**). Importantly, Mulhair et al (2022)^28^ recovered Xenacoela without recoding in reanalyses of the Cannon et al (2016)^18^ and Philippe et al (2019)^26^ datasets when only genes with the greatest ability to recover accepted clades at the gene tree level were applied (**Fig. 1D**).

This previously unacknowledged recovery of Xenacoela across multiple studies, particularly when attempting to minimize phylogenetic error, indicates that this novel clade warrants consideration as an alternative to Acoelomorpha.

### Filtering phylogenomic datasets to reduce error and target internal xenacoelomorph relationships

To better understand the source of the signal for Xenacoela compared to Acoelomorpha I reanalysed datasets from past studies, but with the specific interest of minimising phylogenetic error in resolving the relationships between the three major xenacoelomorph lineages (**Fig. 1E**). I first selected genes that had at least one representative from each of *Xenoturbella*, Acoela and Nemertodermatida to target genes with a basic capability of resolving their relationships (**Fig. 1E**). I then followed the ClanCheck approach^41^, which Mulhair et al (2022) used to filter out genes that perform poorly at recovering generally accepted major clades across the backbone of the animal phylogeny^28^. However, I specifically sought to identify genes where Xenacoelomorpha could be recovered as a clan (the equivalent of a ‘monophyletic’ group in an unrooted tree^47^) in gene trees (**Fig. 1E**). This allowed filtering out of genes where Xenacoelomorpha is not recovered due to *i)* ancient paralogy (where failure of Xenacoelomorpha sequences to form a clan is accurate), or *ii)* limited or biased phylogenetic signal (where failure of Xenacoelomorpha sequences to form a clan is inaccurate, e.g., due to data and/or modelling deficiencies). AU topology tests^48^ indicate that this filtering process enriches for genes that do not reject Xenacoela (**Fig. S1**), suggesting that at least some support for Acoelomorpha is associated with genes that do not recover Xenacoelomorpha.

Bilaterian relationships, including those of Xenacoelomorpha, have previously been shown to be affected by inadequate phylogenetic modelling and systematic errors^26–29,31^. As distant outgroups can worsen these issues and produce incorrect topologies^49,50^, I removed outgroups more distant than Cnidaria, and subsampled off-target bilaterian species to balance outgroup lineage representation and to include only slowly evolving taxa with low levels of missing data (**Fig. 1E**). To further reduce the propensity for systematic error I then used BMGE^51^ to trim alignment sites that either contribute to across taxa compositional heterogeneity or have unusually high variability (**Fig. 1E**).

This approach was applied to three key datasets: the 336 gene ‘best sampled taxa’ dataset from Cannon et al. (2016)^18^, the main 1173 gene dataset from Philippe et al. (2019)^26^, and the least saturated genes dataset used in the main analyses of Marlétaz et al. (2019)^27^. Following filtering this resulted in the following new datasets: Cannon-X (‘X’ for Xenacoelomorpha) with 29 genes, 7448 sites and 28 taxa, Philippe-X with 89 genes, 20847 sites and 29 taxa, and Marlétaz-X with 23 genes, 6627 sites and 28 taxa (**Fig. 1E**). Although these datasets are notably smaller than those in the original studies, they are comparable in size to those recently used by Mulhair et al (2022) to reassess Xenacoelomorpha’s placement in the bilaterian tree of life^28^, and should have a substantially improved signal-to-noise ratio with respect to xenacoelomorph relationships.

### Better modelling of site-heterogeneity reveals support for Xenacoela

Compositional heterogeneity across sites and taxa has been identified as a major source of systematic error contributing to incongruence in animal and bilaterian phylogenomics^26,27,37^. To test absolute model fit in the context of compositional heterogeneity for all three datasets, I employed posterior predictive analyses (PPAs) in Phylobayes^33,37,52^. Specifically I performed the PPA-MEAN and PPA-MAX test of compositional heterogeneity across taxa and the PPA-DIV test of per site amino acid diversity (i.e., compositional heterogeneity across sites)^33,37,52^. These tests were applied to three different modelling strategies: *i)* the standard site-homogeneous LG+G model, *ii)* the site-heterogeneous CAT-GTR+G model, and *iii)* CAT-GTR+G on Dayhoff6 recoded datasets.

In line with the removal of sites associated with compositional heterogeneity across taxa, all datasets passed (at 2 > Z > –2) the PPA-MAX test under all modelling approaches (**Fig. 2A**). However, only Philippe-X passed PPA-MEAN at the amino acid level (**Fig. 2A**), while Cannon336-X fails even when recoded. All datasets drastically fail the PPA-DIV test when the site heterogeneous LG+G model is used, but only Cannon336-X consistently fails, and then barely, under the site-heterogeneous CATGTR+G model (**Fig. 2A**). These results indicate that compositional heterogeneity can be modelled reasonably well for the Philippe-X and Marlétaz-X datasets.

**Figure 2.**
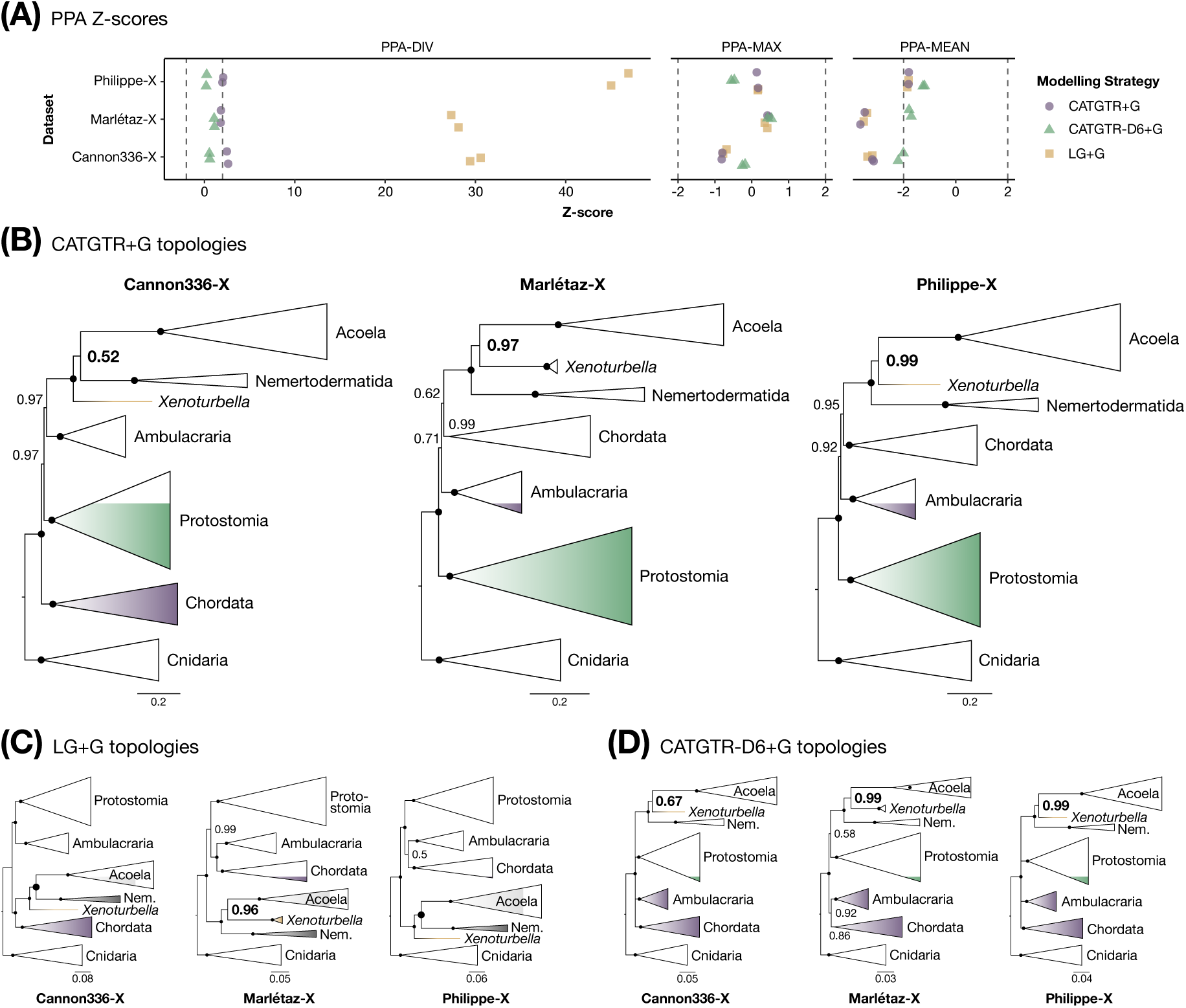
Phylobayes Bayesian analyses of the Cannon336-X, Marlétaz-X and Philippe-X datasets. (**A**) PPA test Z-scores for each dataset and modelling strategy for the PPA-DIV, PPA-MAX, and PPA-MEAN tests, results from each of two Phylobayes chains are shown separately. Dashed vertical lines are positioned at |Z|=2 and |Z|=-2 to indicate pass/fail interpretation. **(B)** CATGTR+G, **(C)** LG+G, and **(D)** CATGTR+D6 topologies for each dataset are shown with species collapsed into major lineages. Posterior probabilities are shown for each node, except for maximal values which are marked by a black circle. Larger font or circle size indicates the support for either the Acoelomorpha or Xenacoela topology. Branch length scale bars represent substitutions/site.

Analysing all three datasets under CATGTR+G always recovers maximal support for the monophyly of Xenacoelomorpha, Acoela, and Nemertodermatida (**Fig. 2B**). However, Acoelomorpha is only recovered when analysing the Cannon336-X dataset, which offers the greatest modelling challenge, and only with equivocal support (Posterior Probability [PP]=0.52; **Fig. 2B**). Instead, I find significant support for Xenacoela, the newly proposed clade with *Xenoturbella* sister to Acoela, in the Philippe-X (PP=0.99) and Marlétaz-X (PP=0.97) analyses. (**Fig. 2B**). By comparison analyses under the less well fitting site-homogeneous LG+G model recover Acoelomorpha with maximal support for the Cannon336-X and Philippe-X analyses, but recover Xenacoela with strong support (PP=0.96) in the Marlétaz-X analysis (**Fig. 3C**). Lastly, although the use of recoding is under debate, Dayhoff6 recoding combined with CATGTR+G offers the best modelling of compositional heterogeneity and recovers Xenacoela for all three datasets, with significant support for Marlétaz-X (PP=0.99) and Philippe-X (PP=0.99) and weak support for Cannon336-X (PP=0.67).

**Figure 3.**
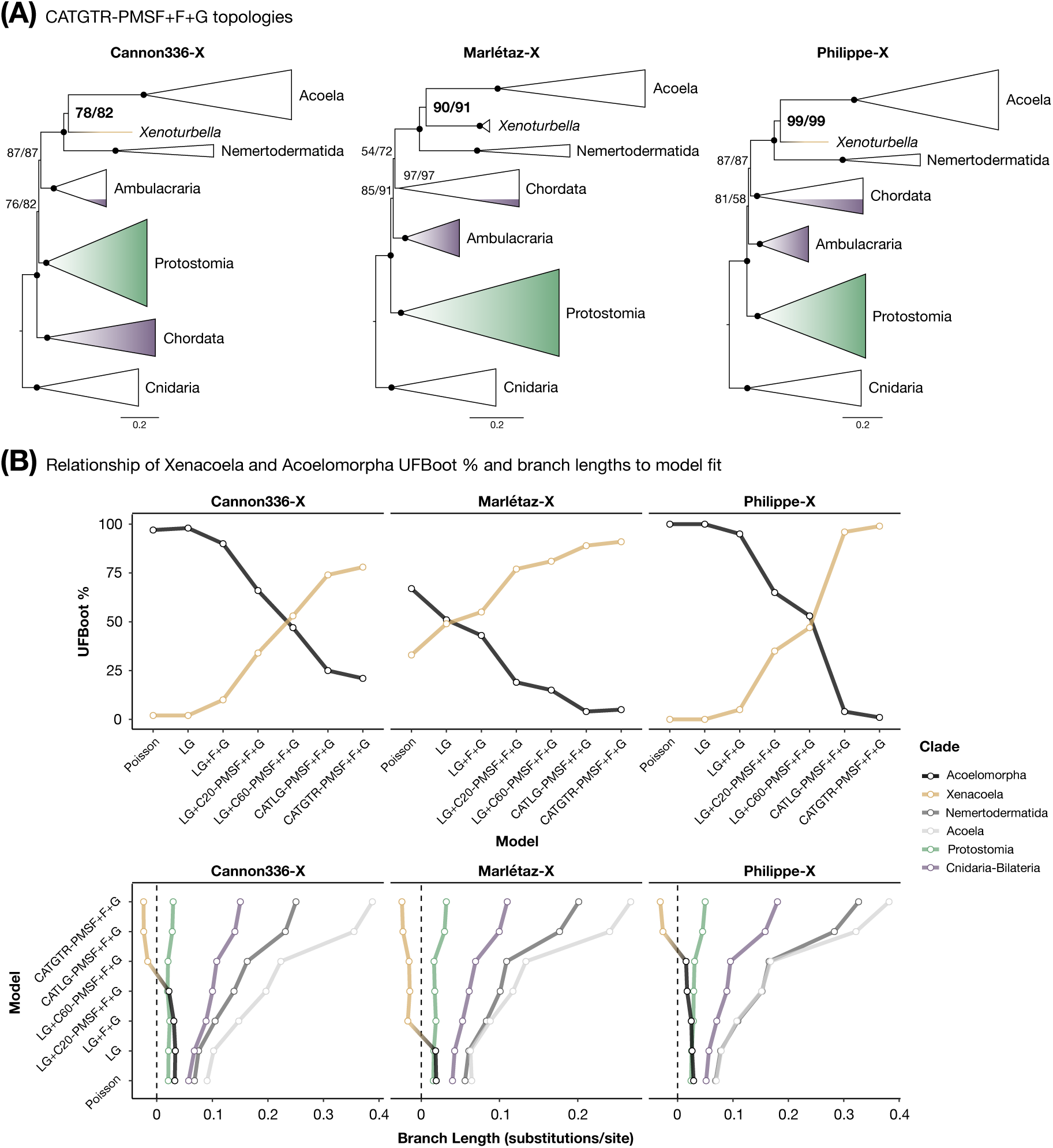
IQ-TREE maximum likelihood analyses of the Cannon336-X, Marlétaz-X and Philippe-X datasets. (**A**) Inferred phylogenies under the best fitting CATGTR-PMSF+F+G (exchangeability matrix and site-specific frequencies inferred by Phylobayes) model for each dataset are shown with species collapsed into major lineages. UFBoot percentages are shown for each node (two values are present as I calculated an exchangeability matrix and site-specific frequencies separately for the two Phylobayes chains and employed both in IQ-TREE analyses), except for maximal values which are marked by a black circle. Larger font indicates the support for either the Acoelomorpha or Xenacoela topology. Branch length scale bars represent substitutions/site. **(B)** UFBoot support values (top) and branch lengths (bottom) for Acoelomorpha and Xenacoela as increasingly more complex and better fitting models are applied (i.e., from Poisson to CATGTR-PMSF+F+G). For branch lengths (bottom) the ancestral Acoelomorpha and Xenacoela branches are treated as the same variable (by treating Acoelomorpha branch lengths as positive and Xenacoela branch lengths as negative) and plotted as a single line for comparison to how model fit alters the length of ancestral branches of other clades.

These results suggest that Acoelomorpha may be an artefact of poorly modelling compositional heterogeneity across sites, with Xenacoela being recovered instead of Acoelomorpha when this is best accounted for.

### Improved model fit correlates with support for Xenacoela and is inversely related to support for Acoelomorpha

To complement the model adequacy analyses in Phylobayes I also performed analyses in IQ-TREE based on relative fit comparisons. The assumption underlying these analyses is that as model fit increases, systematic errors and long branch attraction should be better attenuated. When using the best-fitting (under BIC and AIC) modelling approach tested, the CATGTR-PMSF approach^53^ (which employs site-specific amino acid frequencies and dataset specific amino acid exchangeabilities inferred from Phylobayes) the Xenacoela topology is recovered for all three datasets (ultrafast bootstrap percentage [UFBoot]: Cannon336-X=78/82 [two values are based on separate exchangeability matrix and site specific frequencies inferred from two Phylobayes chains], Marlétaz-X=90/91, Philippe-X=99/99; **Fig. 3A**). On the other hand, when applying the simple, unrealistic, and poorly-fitting Poisson model Acoelomorpha becomes the best supported topology for all three datasets (**Fig. 3B**). Importantly, given that a single best fitting model may be misleading^30,54^, the results reveal a trend where UFBoot support for Acoelomorpha decreases and support for Xenacoela increases as progressively better fitting models are applied (**Fig. 3B**).

Similarly, despite site-heterogeneous models suppressing long-branch attraction, their improved detection of hidden substitutions produces trees with longer branches and a longer total length (where branch length is measured in substitutions per site)^29,31^. This means that well supported clades will often have a longer ancestral branch joining them to the rest of the tree under such models. In line with this, the branch leading to Protostomia and the branch splitting Cnidaria and Bilateria grow longer as model fit increases (**Fig. 3C**). Consistent with the monophyly of each lineage and better detection of hidden substitutions in long-branching or fast-evolving lineages with better fitting site-heterogeneous models, the branches leading to Acoela and Nemertodermatida becomes far longer as model fit improves (**Fig. 3C**). Contrary to the patterns for the above clades, but consistent with UFBoot supports, the branch leading to Acoelomorpha becomes shorter as model fit increases, and eventually switches to a branch leading to Xenacoela that lengthens with further improvement of model fit (**Fig. 3C**).

These results point towards Acoelomorpha being an artefact of model misspecification, that when corrected for reveals support for Xenacoela.

### Simulations implicate Acoelomorpha but not Xenacoela as an error

Simulation-based approaches have recently been employed to compare the propensity for opposing topologies to derive from phylogenetic error^29,31^. The basis of this approach relies on simulating alignments under two opposing topologies and testing whether there is an asymmetry in topology recovery when the data are analysed, particularly when inadequate models are used^29^. Hypothesising, based on my empirical findings, that Acoelomorpha would be easily recovered when correct, and that it might also be recovered by long-branch attraction when incorrect, I performed a number of simulation experiments.

First, taking the basic LG+F+G tree topologies for each datasets, as well as a modification of this topology to produce a tree with the alternative Acoelomorpha or Xenacoela topology, I estimated branch lengths for both trees under the LG+C60-PMSF model in IQ-TREE and then simulated 100 alignments of 25000 sites under the LG+C60-PMSF model for each topology. Analysing these alignments under the simpler LG+F+G model, which should accentuate potential for systematic error^29^, I always recovered the correct topologies except for a small proportion of Cannon336-X simulations recovering Acoelomorpha when Xenacoela is true (**Figure 4A**). This provides little evidence for a bias towards either topology.

**Figure 4.**
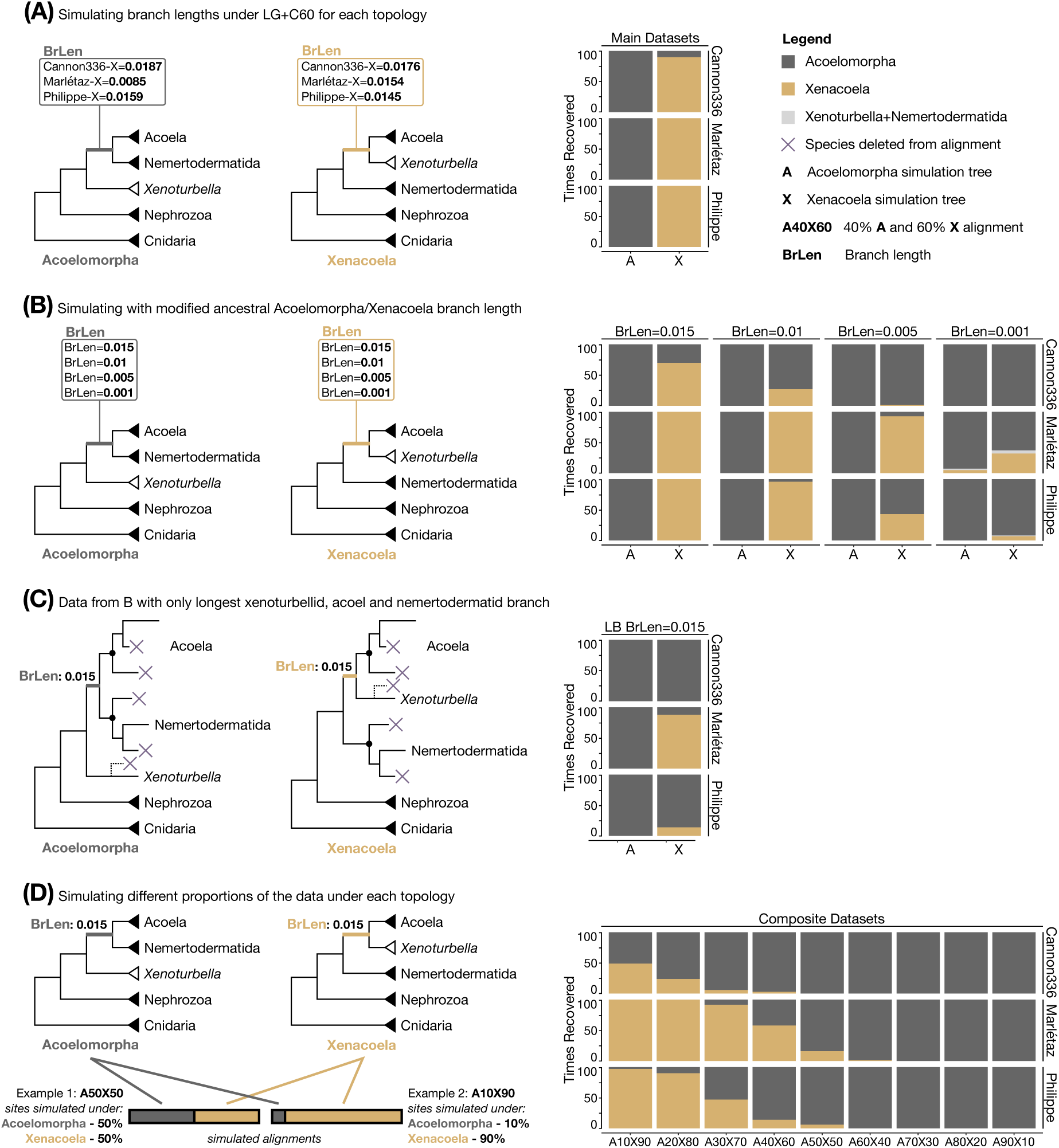
Contrasting recovery frequency of Acoelomorpha and Xenacoela in systematic error inducing simulations. (**A**) Simulations under both the LG+F+G topology and a modified topology supporting the alternative Xenacoela/Acoelomorpha topology for the Cannon336-X, Marlétaz-X, and Philippe-X datasets with branch lengths inferred under LG+C60-PMSF. The inferred empirical branch length supporting each topology is shown for each dataset as well as bar plots recording the number of times each topology is recovered from the 100 replicates of each simulation condition when analysed under LG+F+G (which is simpler than the generating LG+C60-PMSF model). (**B)** Simulations and bar plots as per part (A) but with modified branch lengths to equalise the length of the ancestral branches supporting Acoelomorpha or Xenacoela and assess the influence that the length of this branch has on recovery. (**C**) Bar plot results for the part (**B**) alignment simulations but with the longest (BrLen=0.015) ancestral Acoelomorpha/Xenacoela branch (i.e. most resistant to error) when all but the longest branch species from *Xenoturbella* (only relevant for Marlétaz-X dataset), Acoela and Nemertodermatida are excluded. (**D**) Simulations on the trees from part (**B**) with the longest ancestral Acoelomorpha/Xenacoela branch (BrLen=0.015) but with different proportions of the alignment simulated under each tree topology rather than all under a single topology, as well as a bar plots of topology recovery count. Nephrozoa is shown as a clade to simplify presentation (although it is present in many of the LG+F+G fixed topologies used for simulation). The *Xenoturbella* clade is shown in white in parts (**A)**, (**B**), and (**D**), while a dotted line branch is used to show removed *Xenoturbella* species in part (**C**), as only Marlétaz-X contain more than one xenoturbellid.

To better explore how the length of the ancestral Acoelomorpha/Xenacoela branch influences recovery of the simulating tree I next modified the main LG+C60-PMSF branch length trees to remove the ancestral branch length asymmetry between these topologies (**Figure 4B**). I first altered the ancestral Acoelomorph/Xenacoela branch length of each topology to 0.015 substitutions/site. Simulating and Analysing inferred trees produces results consistent with the main simulations where the correct topology was always recovered, except for sometimes recovering Acoelomorpha when Xenacoela is correct for the Cannon336-X analyses (**Figure 4B**). I then tested gradual reduction of the equalised ancestral branch lengths to 0.01, 0.005, and 0.001 substitutions/site (**Figure 4B**). This revealed a very clear pattern that cannot be explained by differences in the simulation tree. Acoelomorpha is almost always recovered when correct, even when the ancestral Acoelomorpha branch is very short (**Figure 4B**). The only exception to this is at the shortest ancestral Acoelomorpha branch length for Marlétaz-X, where Acoelomorpha recovery drops slightly to 93% of cases. On the other hand, Xenacoela recovery is highly sensitive to ancestral branch length, with Acoelomorpha being erroneously recovered in 63-100% of cases where the ancestral Xenacoela branch is shortest (**Figure 4B**).

To complement this, I also reanalysed the 0.015 substitutions/site ancestral branch alignments but this time removed all but the fastest evolving acoel, nemertodermatid and, in the case of the Marlétaz dataset (which has more than one xenoturbellid), xenoturbellid, as this should increase the potential for long-branch attraction errors^31^ (**Figure 4C**). These data revealed that Acoelomorpha was always recovered when correct, and was also often erroneously recovered, although at very different frequencies across simulating data (Cannon336-X: 100%, Marlétaz-X: 12%, Philippe-X: 86%), when Xenacoela was correct (**Figure 4C**).

To help understand the influence that orthology errors (and other factors such as incomplete lineage sorting or gene flow) might have on the recovery of Acoelomorpha or Xenacoela I reperformed simulations using the modified topologies with 0.015 substitutions/site ancestral Acoelomorpha or Xenacoela branches. However, this time varying proportions of the data were simulated under each tree^29^ (**Figure 4D**). In total I tested nine data composition variants spanning 10% windows from 10% Acoelomorpha and 90% Xenacoela to 90% Acoelomorpha and 10% Xenacoela (**Figure 4D**). If unbiased the data might be expected to produce either topology roughly 50% of the time when 50% of the data is simulated under each topology^29^. However, I observe a clear bias in favour of Acoelomorpha, which is always recovered (with the exception of a single tree with 60% of the data simulated under Acoelomorpha) when it is the majority simulation tree in the data and is always the most frequently recovered topology when 50% of the data are simulated under each topology (**Figure 4D**). Conversely, even when 90% of the data are simulated under Xenacoela, Acoelomorpha is still recovered in a small number of cases for the Philippe-X analyses, and in the majority of cases for the Cannon336-X analyses (**Figure 4D**).

These simulations highlight the influence of branch length, which is an important factor influencing branching order inference in animal phylogenomics^29,31,55,56^, on simulation outcomes, and clearly indicate that Acoelomorpha is not only far more easily recovered than Xenacoela when correct, but can also easily be recovered in error in place of Xenacoela.

### Support for Nephrozoa is inversely related to model fit

Beyond internal Xenacoelomorph relationships, accurately placing Xenacoelomorpha in the bilaterian tree of life is among the most vexing problems in animal phylogenomics^18,26,28,29^. While the datasets used here are small and internal Xenacoelomorpha-targeted, the analyses reveal a clear pattern of support for Nephrozoa being suppressed as better fitting models are applied (**Fig. 2A-C and Fig. 5**), consistent with most recent reports^26–30^. However, despite Xenambulacraria being the primary alternative hypothesis^3,26–28^, it is only recovered for the Cannon336-X dataset (**Fig. 2B**, **Fig. 3A**), while unexpected support emerges for Xenacoelomorpha as sister to Chordata for the Marlétaz-X and Philippe-X datasets (**Fig. 2B**, **Fig. 3A**). However, this support is ameliorated when the data are Dayhoff6 recoded (**Fig. 2D**). While these findings do not point towards a consistently well supported sister group to Xenacoelomorpha, they do reveal an inverse relationship between model fit and support for Nephrozoa, indicating that it is likely a systematic error.

**Figure 5.**
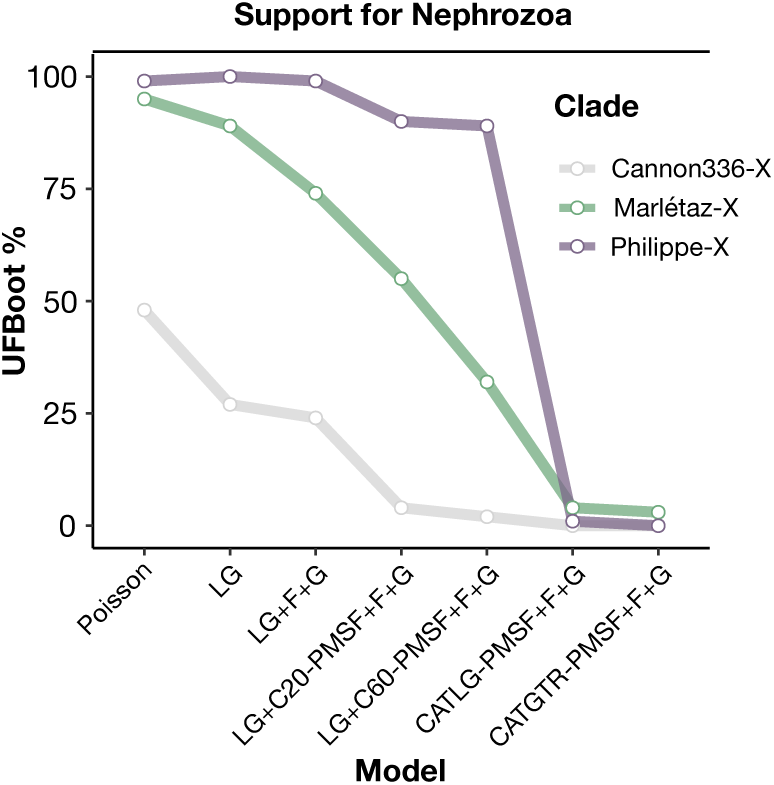
Support values for Nephrozoa in IQ-TREE maximum likelihood analyses of the Cannon336-X, Marlétaz-X and Philippe-X datasets. UFBoot % support values favouring the Nephrozoa hypothesis as increasingly more complex and better fitting models are applied (i.e., from Poisson to CATGTR-PMSF+F+G).

## Discussion

The phylogenetic placement of the three xenacoelomorph lineages has been a long standing problem in evolutionary biology^4,18,21,28,57^, with *Xenoturbella* having been described as the ‘champion wanderer’ of bilaterian phylogeny^21^. This study sets *Xenoturbella* on its way once more, nesting it deeper within Xenacoelomorpha as the sister group to Acoela. The proposed name for this clade, Xenacoela, is consistent with the naming of Xenacoelomorpha^3^, and Xenambulacraria^12^, and does not rely on interpretations of morphological character history. I suggest retaining Xenacoelomorpha as the phylum name, rather than simply including Xenoturbellida within Acoelomorpha, as this will maintain coherence with other proposed clade names, such as Xenambulacraria^12^. I propose that Xenacoela should take the place of Acoelomorpha, which appears to be invalid based on my findings, as a subphylum to Xenacoelomorpha. If accepted, this would also require additional taxonomic revisions; for example, Xenoturbellida might be demoted to class alongside Acoela, and Nemertodermatida raised as the other xenacoelomorph subphylum.

My results break the monophyly of the acoelomorph flatworms, which I propose derives from long branch attraction between the fast-evolving Acoela and Nemertodermatida. The remarkably long branches of Acoela raise the possibility that further suppression of long-branch attraction could recover a Xenacoela variant where *Xenoturbella* falls within, instead of sister to, Acoela. However, this seems unlikely, as while the consistent recovery of maximal support across analyses for the monophyly of Acoela could in theory be driven by a strong systematic error signal, branch length analyses reveal that the branch leading to Acoela becomes dramatically longer as model fit improves (**Fig. 3B**). This lends confidence in the placement of *Xenoturbella* as sister to, and not within, Acoela.

The monophyly of Acoelomorpha was also questioned in the pre-phylogenomic era^24,58–60^. However, this predated discovery of Xenacoelomorpha, and specifically referred to Nemertodermatida being sister to Nephrozoa, with Acoela sister to both at the root of the bilaterian tree, a topology that supported an acoelomorph flatworm-like bilaterian ancestor^24,58,59^. Although I also propose that Acoelomorpha is not monophyletic, my findings are distinct from past inferences, as i) the analyses here recover Xenacoelomorpha (albeit unsurprising given that only genes that recover Xenacoelomorpha as a clan at the gene tree level were used), ii) the non-monophyly of Acoelomorpha is with respect to *Xenoturbella* rather than Nephrozoa, and iii) consistent with most recent studies^26–30^, my analyses indicate that Nephrozoa is likely a systematic error—ameliorating the phylogenetic evidence for an acoelomorph-like last common bilaterian ancestor.

The evidence here that Acoela and Nemertodermatida cause long-branch attraction even within Xenacoelomorpha provides an auxiliary line of support to past arguments that Xenacoelomorpha falling as sister to Nephrozoa is a long-branch attraction artefact^3,26,29^. It also lends credence to past analyses using only the slower evolving *Xenoturbella* as representative for all of Xenacoelomorpha, which has previously shifted support away from Nephrozoa and towards Xenambulacraria when site-heterogeneous models were employed^26^. My analyses also find that support for Nephrozoa is associated with simple, poorly fitting models, in line with recent studies^26–28^. However, I do not recover a strong and consistent signal for any single closest relative to Xenacoelomorpha across datasets and analyses when better fitting models are used. A link with deuterostomes, as sister to Ambulacraria (i.e., Xenambulacraria, sensu^3,26–28^) or Chordata (no previous phylogenomic evidence), seems most likely based on amino acid analyses (**Figs. 2B and 3A**). However, monophyletic deuterostomes are not always recovered (**Figs. 2B and 3A**), a possibility that has been raised in recent studies^26,31^, and the support for this deuterostome affinity is attenuated with Dayhoff6 recoding (**Fig. 2D**). Importantly, the datasets used here are dramatically reduced compared to those used in the original studies^18,26,27^, and only include genes recovering Xenacoelomorpha as a clan at the gene tree level. While using these genes should produce datasets with more coherent signal for the relationship between Xenacoelomorpha and other animals, it is not clear that this is the case, as I did not also consider the recovery of expected clans beyond Xenacoelomorpha as done by Mulhair et al (2019)^28^.

In this context, many questions about gene choice, dataset trimming, phylogenetic modelling approaches, amino acid recoding and their intersection remain open and debated in phylogenomics^28,30,31,35,41,53,61–70^. Nonetheless, I am optimistic that strategies such as that applied here can provide a path forward for future efforts to detect and resolve previously hidden long-branch attraction artefacts in the tree of life. The stringent, but focused, data filtering approach, when paired with well-fitting models, appears to minimize branching artefacts by improving signal-to-noise ratio, and dramatically reduces computational requirements, making phylogenomic analyses more environmentally friendly^71^ and reproducible. While this comes at notable loss of data, it also appears that at least some such data, with available modelling and orthology inference approaches, may be misleading.

Importantly, Xenacoela recovery is not restricted to a single set of stringent conditions. Instead, the topology can be recovered using multiple datasets and approaches thought to reduce phylogenetic error. For example, different subsampling of genes^26,28^, or the combined application of site-heterogeneous models and recoding to large phylogenomic datasets of over 1000 genes^26^, have both recovered Xenacoela (**Fig. 1D**). It is also noteworthy that Xenacoela joins the shortest and longest branching Xenacoelomorph lineages, instead of the two longest branching lineages (i.e. Acoelomorpha) as might be expected in the case of common errors like long-branch attraction. The simulations performed here lend further support to this theory, showing that Acoelomorpha is almost always recovered when it is the correct tree and is also easily recovered in error. Meanwhile, Xenacoela is more difficult to recover when correct and is only recovered in error in very rare edge cases. These simulations also indicate the importance of careful simulation set up and consideration of different starting datasets to enable fair and general interpretation of results.

In summary, my results reject Acoelomorpha in favour of Xenacoela, and indicate that Nephrozoa is likely to be a systematic error. Development of chromosome-scale genome sequences from across Xenacoelomorpha will allow further comparison of support for Xenacoela or Acoelomorpha, with syntenic evidence producing larger and more reliable ortholog sets and potentially providing additional phylogenetic characters in the form of rare genome changes^72–74^. Additionally, I predict that careful taxonomic consideration of Xenacoela will reveal morphological characters that represent synapomorphies uniting *Xenoturbella* and Acoela, as a formal reappraisal of support for Acoelomorpha has not been performed since establishment of Xenacoelomorpha and very few morphological characters are known to unite Acoelomorpha^75^.

## Methods

### Dataset Preparation

I employed three datasets derived from previous studies focused on placing Xenacoelomorpha amongst Bilateria and filtered these to hone in on internal Xenacoelomorpha relationships and limit propensity for systematic errors. The first of these was the secondary 336 gene dataset (56 taxa, 81451 sites, 11% missing data) from Cannon et al (2016)^18^, which recovered Nephrozoa in the original study. This dataset includes fewer taxa with lower levels of missing data and more genes than the main 212 gene dataset (78 taxa, 44896 sites, 31% missing data) employed by Cannon et al (2016)^18^, and was generated using the same dataset assembly protocol (orthology inference, alignment, trimming, paralog pruning etc.). I employed the 336 gene dataset for analyses as I judged a preliminary filtered dataset produced from the 212 gene dataset as too small (**Fig. S2A**). I also reused the main 1173 gene dataset (59 taxa, 350088 sites, 23.5% missing data) from Philippe et al. (2019)^26^, which supported Xenambulacraria in the original study, is the largest dataset yet to be applied to Xenacoelomorpha, relies on genomes as well as transcriptome data for xenacoelomorphs, and reportedly contains fewer data errors than other datasets. Lastly, I considered the main least saturated dataset of Marlétaz et al. (2019)^27^, which although not focused specifically on Xenacoelomorpha’s placement among Bilateria, is the only phylogenomic dataset to include two *Xenoturbella* species alongside representatives of both Acoela and Nemertodermatida. I specifically relied on the ‘Broad’ dataset including only the least saturated genes (258 genes, 103 taxa, 74014 sites, 29.16% missing data), as used in the analyses in the original study that employed site-heterogeneous models, and supported Xenambulacraria^27^. I also attempted to employ the 422 gene pan-Metazoan dataset of Laumer et al. (2019) but this produced a relatively short alignment after filtering which appeared to have weak resolving power for the problem of interest (e.g. adjacent nodes to Acoelomorpha/Xenacoela in the tree, including Xenacoelomorpha and Nemertodermatida were weakly supported) and so these data were not analysed further (**Figs. S2A and S2B**).

I filtered each of these datasets by gene to select genes with a greater potential to accurately infer internal xenacoelomorph relationships. To do this I first selected only genes for which there was at least one sequence representative from each of *Xenoturbella*, Acoela, and Nemertodermatida present, a basic requirement to resolve the relationships between these lineages. In addition, I only retained genes in which Xenacoelomorpha sequences form a clan^47^ (i.e., the Xenacoelomorpha sequences are monophyletic assuming the tree root falls outside Xenacoelomorpha in the gene tree) at the gene tree level. The assumption of this step is that genes where Xenacoelomorpha forms a clan should be better enriched for either orthology and/or phylogenetic signal, as failure to recover Xenacoelomorpha in a gene tree most likely derives from either ancient paralogy between xenacoelomorph sequences or poor signal for the clade. To test this I used the gene trees inferred by Mulhair et al. (2022)^28^ for the main 212 gene Cannon dataset (which as explained above I did include in later analyses; from the ‘cannon_2016/OG_data/OG_trees/all_trees’ directory of https://github.com/PeterMulhair/Xenaceol_Paralogy^28^) and the Philippe (2019) dataset (from the ‘phillipe_2019/OG_data/OG_trees/all_trees’ directory of https://github.com/PeterMulhair/Xenaceol_Paralogy^28^). For the Cannon336 dataset I extracted individual gene alignments from the original supermatrix alignment (hamstr_best_coverage_taxa.phy from https://datadryad.org/stash/dataset/doi:10.5061/dryad.493b7^18^) using the associated partition coordinates (“README_for_hamstr_best_coverage_taxa.txt” from https://datadryad.org/stash/dataset/doi:10.5061/dryad.493b7^18^) for each gene alignment using the ‘split_supermatrix_to_genes.py’ script from https://github.com/wrf/supermatrix^65^, and removed species for which there was no sequence data for that partition using BMGE (version 1.12; flags: –h 1 –g 1)^51^. For the Marlétaz et al. (2019)^27^ data I compared trimmed individual gene alignments (in ‘alis_filtered.tgz’ from https://zenodo.org/record/1403005^27^) to the ‘Broad’ least saturated genes supermatrix (‘Concat-Tc111217-broad.phy’ in ‘Concat-alis.tgz’ from https://zenodo.org/record/1403005^27^) and retained those gene alignments that were contained within (as a subset of) the supermatrix for further analysis. I then performed phylogenetic analysis on the genes of Cannon et al. (2016)^18^ 336 gene dataset and the Marlétaz et al. (2019)^27^ genes following the approach applied by Mulhair et al^28^ on the 212 gene Cannon et al. (2016)^18^ dataset and the Philippe et al. (2019)^26^ dataset. Briefly, I used IQ-TREE (version 1.6.12)^76^, specifying 1000 ultrafast bootstraps (-bb 1000)^77^ and for ModelFinder^78^ to select the best-fitting model (-m TEST). At this point gene trees from each dataset were analysed to test whether Xenacoelomorpha formed a clan using ClanCheck (https://github.com/ChrisCreevey/clan_check)^41^, and those where it did were retained and combined into supermatrices for each dataset.

I next filtered species and sites from these datasets. As distant outgroups can mislead phylogenetic analyses^49,50^, I removed species more distantly related to Bilateria than Cnidaria, and also subsampled outgroup bilaterian lineages to remove fast-evolving or missing data replete off-target species, as well as to balance taxon sampling between major outgroup lineages (this point represents the X-noTrim ‘error prone’ datasets). As compositional heterogeneity and fast and variable evolutionary rates are important factors influencing bilaterian phylogeny^26,27,31^, I stripped sites from each supermatrix to reduce compositional heterogeneity across taxa and unexpectedly high variability using BMGE (version 1.12; flags: –m BLOSUM95 –s FAST)^51^.

The above filtering steps resulted in three new datasets used for them main analysis: Cannon-X with 29 genes, 7448 sites and 28 taxa, Philippe-X with 89 genes, 20847 sites and 29 taxa, and Marlétaz-X with 23 genes, 6627 sites and 28 taxa.

To understand how using only genes that recover Xenacoelomorpha might relate to support for Aceolomorpha or Xenacoela, AU topology test comparisons^48^ were performed in IQ-TREE (version 1.6.12)^76^ using 10000 RELL replicates^79^ on each gene family in each of the three main ‘-X’ datasets to assess gene level support for Xenacoela in comparison to Acoelomorpha. Genes were analysed after subsampling species but prior to trimming sites (which was performed as a single step on the concatenated supermatrices as suggested by BMGE). The LG+G+F topology was used for each gene family along with a modification of this tree to allow testing of the remaining two topologies from Acoelomorpha, Xenacoela, and Xenoturbellida+Nemertodermatida. Each tree topology was pruned to match the species present in each gene alignment using a custom python script employing the ETE3 toolkit^86^. For comparison to this dataset the same analysis was performed on the species subsampled alignments for those genes which did not recover Xenacoelomorpha (but still had at least one representative from each of Acoela, Nemertodermatida and Xenoturbellida).

Although not included in the main analyses, multiple variant phylogenetic analyses were performed without trimming taxa or without trimming compositionally biased or highly variable sites to better understand the influence of each of these filtering steps. IQ-TREE analysis of these datasets (as described below) under precomputed site-homogeneous and site-heterogeneous models generally revealed a similar pattern of support moving from Acoelomorpha and towards Xenacoela (**Fig. S3**). Unsurprisingly this trend was more subdued without or with less stringent site-stripping (**Fig. S3**). Site-stripping on datasets without species subsampling produced shorter alignments (<1000 positions for Marlétaz), but revealed support for Xenacoela for the Philippe dataset even without site-heterogeneous models (**Fig. S3, Fig. S4**).

### Bayesian posterior predictive analyses and phylogenetics

I used Phylobayes (version 4.1c) to perform Bayesian phylogenetic analyses and posterior predictive simulation analyses (PPA)^32,33,37,52^. I ran two Markov Chain Monte Carlo chains for 10000 points each using the ‘pb’ command under LG+G^80,81^, CATGTR+G^32^, and Dayhoff6^34^ recoded under CATGTR+G for each filtered dataset. Convergence was assessed and consensus trees produced for each modelling approach for each dataset using the ‘bpcomp’ command, with the first 5000 points (50% of each chain) discarded as burn-in and requiring the ‘maxdiff’ value to be less than 0.3^52^. PPAs^33,37^ were performed using the ‘ppred’ command, and the same 5000 point burn-in as for ‘bpcomp’. The ‘-sat’ flag was specified to perform the PPA-DIV (average per-site amino acid diversity) analyses and the ‘-comp’ flag used to perform the PPA-MAX and PPA-MEAN (maximum and average compositional heterogeneity across taxa) analyses and produce associated z-scores^33,52^. The ‘readpb’ command^52^ was used separately for every chain to extract the inferred site-specific amino acid frequencies (‘-ss’ flag), and the mean inferred amino acid exchangeability matrix (‘-rr’ flag).

### Maximum likelihood phylogenetics and relative model fit comparisons

IQ-TREE^76^ (version 1.6.12), with 1000 ultrafast bootstrap replicates (-bb 1000)^77^ was used for all maximum likelihood phylogenetic analyses, and relative model fit comparisons were based on the Akaike Information Criterion (AIC) and the Bayesian Information Criterion (BIC) values inferred using IQ-TREE’s built in ModelFinder tool^78^. I employed a suite of models with very different properties to analyse support for Acoelomorpha and Xenacoela in comparison to relative model fit, which has the benefit of not relying upon the topology and support value derived from a single best-fit model alone^30,54^. This spanned i) the simple, equal exchangeabilities and frequencies of the Poisson model (Poisson), ii) the more realistic exchangeabilities of the LG model (LG), iii) combining LG exchangeabilities^80^ with 4 discrete gamma categories for rate heterogeneity^81^ as well as empirical amino acid frequencies from the data (LG+F+G), iv) combining iii with the precomputed site-heterogeneous C20^82^ frequencies model, with posterior mean site frequencies (PMSF)^83^ inferred under consensus tree from iii (LG+C20-PMSF+F+G), v) As per iv but with C60^82^ instead of C20 frequency model (LG+C60-PMSF+F+G), vi) as per iii but using the site-specific amino acid frequencies inferred in Phylobayes (CATLG-PMSF+F+G), vii) as per vi but also using the mean amino acid exchangeability matrix inferred in Phylobayes instead of the LG model (CATGTR-PMSF+F+G). Two analyses each were ran for vi and vii as two Phylobayes chains were used separately to generate site-specific amino acid frequencies and a mean amino acid exchangeability matrix (bootstrap results for both are shown in Figure 1A, but only those with the best AIC and BIC values are plotted in Figure 1B and Figure S1G). These Phylobayes exchangeability matrices and site frequencies were converted to IQ-TREE format using the CAT-PMSF ‘convert-exchangeabilities.py’ and ‘convert-site-dists.py’ scripts from https://github.com/drenal/cat-pmsf-paper/tree/main/scripts^53^. Models vi and vii follow the CAT-PMSF approach^53^ except for not using a fixed topology for Phylobayes inference of site frequencies and amino acid exchangeabilities. Branch length and bootstrap values for branches/clades of interest were manually extracted from the IQ-TREE consensus trees.

### Systematic error simulations analyses

I set up simulation experiments to compare the accurate and inaccurate recovery of Acoelomorpha and Xenacoela under systematic error conditions^29,31^. To do so I first took the basic LG+F+G IQ-TREE tree topologies generated with each dataset (recovering Acoelomorpha for Cannon336-X and Philippe-X and Xenacoela for Marlétaz-X), and also modified the branching order to have an alternative tree (to Xenacoela for Cannon336-X and Philippe-X and to Acoelomorpha for Marlétaz-X). I then used each fixed topology to estimate branch lengths under the more complex and better fitting LG+C60-PMSF model in IQ-TREE. For each of these 6 trees we then simulated 100 alignments of 25000 sites using AliSim^84^ in IQ-TREE (version 2.2.0^85^ used for data simulation steps only) under LG+C60-PMSF. To provide systematic error conditions I analysed all of these alignments under the simpler LG+F+G model.

The above analyses suggested an important influence of the length of the branch leading to Acoelomorpha or Xenacoela on the inference and so I next performed simulations and analyses exactly as above but with the length of the branch leading to Acoelomorpha or Xenacoela modified. The aim of this was two-fold: i) to make the comparison fair, such that the branch leading to either clade is of equal length (and so should not influence differences in topology recovery), and ii) to assess the impact that the length of this branch has on recovery of either topology. To do this, I manually edited the length of the branch leading to Acoelomorpha or Xenacoela to be 0.015 substitutions/site in length and also tested gradual reduction of this length by setting alternative values of 0.01, 0.005, and 0.001 substitutions per site for these branch lengths. In all cases I redistributed the difference between the inferred branch length and the modified branch length to/from the immediate daughter branches to maintain the root to tip distances (and total inferred substitutions in some sense) along the tree. I applied this such that if the branch leading to Acoelomorpha or Xenacoela is lengthened or shortened, then the immediate daughter branches are equally shortened or lengthened, respectively. In all cases I simulated 100 alignments of 25000 sites each using AliSim in IQ-TREE under LG+C60-PMSF, and analysed these data under LG+F+G model.

To complement the above I also reanalysed (again under LG+F+G) the simulated datasets with equalised 0.015 substitutions/site branch lengths leading to Acoelomorpha and Xenacoela but this time with only the longest branching of each of the three Xenacoelomorpha lineages included, as this should further increase the potential for long-branch attraction errors.

To better assess the influence that orthology errors (and other factors such as incomplete lineage sorting or gene flow) might have on the recovery of Acoelomorpha or Xenacoela I reperformed simulations under the LG+C60-PMSF branch length trees with modified to 0.015 substitutions/site ancestral Acoelomorpha or Xenacoela branches. However, this time a varying proportion of the data was simulated under each tree topology^29^. In total I tested nine data composition variants spanning 10% windows from 10% Xenacoela and 90% Acoelomorpha to 90% Xenacoela and 10% Acoelomorpha. If unbiased the data might be expected to produce either topology roughly 50% of the time when 50% of the data is simulated under each topology^29^. In all cases I simulated 100 alignments of 25000 sites each using AliSim in IQ-TREE under LG+C60-PMSF, and analysed these data under LG+F+G model.

Custom dataset-specific python scripts using the ETE3 toolkit^86^ were used to parse simulation results and report the xenacoelomorph relationships recovered in all simulation trees.

## Acknowledgments

I thank Aoife McLysaght for comments on an early version of this manuscript and Lénárd L. Szánthó for suggesting use of the CAT-PMSF approach. I am supported by an Irish Research Council Government of Ireland Postdoctoral Fellowship (GOIPD/2021/466).

## Author Contributions

AKR conceived and designed the study, performed analyses, and prepared the manuscript.

## References

1. Jondelius, U., Raikova, O.I., and Martinez, P. (2019). Xenacoelomorpha, a Key Group to Understand Bilaterian Evolution: Morphological and Molecular Perspectives. In Evolution, Origin of Life, Concepts and Methods, P. Pontarotti, ed. (Springer International Publishing), pp. 287–315. 10.1007/978-3-030-30363-1_14.

2. Hejnol, A., and Pang, K. (2016). Xenacoelomorpha’s significance for understanding bilaterian evolution. Current Opinion in Genetics & Development 39, 48–54. 10.1016/j.gde.2016.05.019.

3. Philippe, H., Brinkmann, H., Copley, R.R., Moroz, L.L., Nakano, H., Poustka, A.J., Wallberg, A., Peterson, K.J., and Telford, M.J. (2011). Acoelomorph flatworms are deuterostomes related to Xenoturbella. Nature 470, 255–258. 10.1038/nature09676.

4. Ruiz-Trillo, I., and Paps, J. (2016). Acoelomorpha: earliest branching bilaterians or deuterostomes? Org Divers Evol 16, 391–399. 10.1007/s13127-015-0239-1.

5. Gavilán, B., Perea-Atienza, E., and Martínez, P. (2016). Xenacoelomorpha: a case of independent nervous system centralization? Philosophical Transactions of the Royal Society B: Biological Sciences 371, 20150039. 10.1098/rstb.2015.0039.

6. Martínez, P., Hartenstein, V., and Sprecher, S.G. (2017). Xenacoelomorpha Nervous Systems. In Oxford Research Encyclopedia of Neuroscience 10.1093/acrefore/9780190264086.013.203.

7. Perea-Atienza, E., Gavilán, B., Chiodin, M., Abril, J.F., Hoff, K.J., Poustka, A.J., and Martinez, P. (2015). The nervous system of Xenacoelomorpha: a genomic perspective. Journal of Experimental Biology 218, 618–628. 10.1242/jeb.110379.

8. Andrikou, C., Thiel, D., Ruiz-Santiesteban, J.A., and Hejnol, A. (2019). Active mode of excretion across digestive tissues predates the origin of excretory organs. PLOS Biology 17, e3000408. 10.1371/journal.pbio.3000408.

9. Abalde, S., Tellgren-Roth, C., Heintz, J., Pettersson, O.V., and Jondelius, U. (2023). The draft genome of the microscopic Nemertoderma westbladi sheds light on the evolution of Acoelomorpha genomes. 2023.06.28.546832. 10.1101/2023.06.28.546832.

10. Tyler, S., and Schilling, S. (2011). Phylum Xenacoelomorpha Philippe, et al., 2011. In: Zhang, Z.-Q. (Ed.) Animal biodiversity: An outline of higher-level classification and survey of taxonomic richness. Zootaxa 3148, 24–25. 10.11646/zootaxa.3148.1.6.

11. Ehlers, U. (1985). Das Phylogenetische System der Plathelminthes (Stuttgart, New York: Gustav Fischer Verlag).

12. Bourlat, S.J., Juliusdottir, T., Lowe, C.J., Freeman, R., Aronowicz, J., Kirschner, M., Lander, E.S., Thorndyke, M., Nakano, H., Kohn, A.B., et al. (2006). Deuterostome phylogeny reveals monophyletic chordates and the new phylum Xenoturbellida. Nature 444, 85–88. 10.1038/nature05241.

13. Franzén, Å., and Afzelius, B.A. (1987). The ciliated epidermis of Xenoturbella bocki (Platyhelminthes, Xenoturbellida) with some phylogenetic considerations. Zoologica Scripta 16, 9–17. 10.1111/j.1463-6409.1987.tb00046.x.

14. Lundin, K. (1998). The epidermal ciliary rootlets of Xenoturbella bocki (Xenoturbellida) revisited: new support for a possible kinship with the Acoelomorpha (Platyhelminthes). Zoologica Scripta 27, 263–270. 10.1111/j.1463-6409.1998.tb00440.x.

15. Westblad, E. (1949). Xenoturbella bocki n. g., n. sp. a peculiar, primitive Turbellarian type. Arkiv for Zoologi 1, 3–29.

16. Nakano, H., Lundin, K., Bourlat, S.J., Telford, M.J., Funch, P., Nyengaard, J.R., Obst, M., and Thorndyke, M.C. (2013). Xenoturbella bocki exhibits direct development with similarities to Acoelomorpha. Nat Commun 4, 1537. 10.1038/ncomms2556.

17. Hejnol, A., Obst, M., Stamatakis, A., Ott, M., Rouse, G.W., Edgecombe, G.D., Martinez, P., Baguñà, J., Bailly, X., Jondelius, U., et al. (2009). Assessing the root of bilaterian animals with scalable phylogenomic methods. Proceedings of the Royal Society B: Biological Sciences 276, 4261–4270. 10.1098/rspb.2009.0896.

18. Cannon, J.T., Vellutini, B.C., Smith, J., Ronquist, F., Jondelius, U., and Hejnol, A. (2016). Xenacoelomorpha is the sister group to Nephrozoa. Nature 530, 89–93. 10.1038/nature16520.

19. Nielsen, C. (2010). After all: Xenoturbella is an acoelomorph!: Xenoturbella is an acoelomorph. Evolution & Development 12, 241–243. 10.1111/j.1525-142X.2010.00408.x.

20. Bourlat, S.J., Nielsen, C., Lockyer, A.E., Littlewood, D.T.J., and Telford, M.J. (2003). Xenoturbella is a deuterostome that eats molluscs. Nature 424, 925–928. 10.1038/nature01851.

21. Nakano, H. (2015). What is Xenoturbella? Zoological Letters 1, 22. 10.1186/s40851-015-0018-z.

22. Philippe, H., Brinkmann, H., Martinez, P., Riutort, M., and Baguñà, J. (2007). Acoel Flatworms Are Not Platyhelminthes: Evidence from Phylogenomics. PLOS ONE 2, e717. 10.1371/journal.pone.0000717.

23. Norén, M., and Jondelius, U. (1997). Xenoturbella’s molluscan relatives⃛. Nature 390, 31–32. 10.1038/36242.

24. Wallberg, A., Curini-Galletti, M., Ahmadzadeh, A., and Jondelius, U. (2007). Dismissal of Acoelomorpha: Acoela and Nemertodermatida are separate early bilaterian clades. Zoologica Scripta 36, 509–523.

25. Rouse, G.W., Wilson, N.G., Carvajal, J.I., and Vrijenhoek, R.C. (2016). New deep-sea species of Xenoturbella and the position of Xenacoelomorpha. Nature 530, 94–97. 10.1038/nature16545.

26. Philippe, H., Poustka, A.J., Chiodin, M., Hoff, K.J., Dessimoz, C., Tomiczek, B., Schiffer, P.H., Müller, S., Domman, D., Horn, M., et al. (2019). Mitigating Anticipated Effects of Systematic Errors Supports Sister-Group Relationship between Xenacoelomorpha and Ambulacraria. Curr Biol 29, 1818–1826.e6. 10.1016/j.cub.2019.04.009.

27. Marlétaz, F., Peijnenburg, K.T.C.A., Goto, T., Satoh, N., and Rokhsar, D.S. (2019). A New Spiralian Phylogeny Places the Enigmatic Arrow Worms among Gnathiferans. Current Biology 29, 312–318.e3. 10.1016/j.cub.2018.11.042.

28. Mulhair, P.O., McCarthy, C.G.P., Siu-Ting, K., Creevey, C.J., and O’Connell, M.J. (2022). Filtering artifactual signal increases support for Xenacoelomorpha and Ambulacraria sister relationship in the animal tree of life. Current Biology 32, 5180–5188.e3. 10.1016/j.cub.2022.10.036.

29. Kapli, P., and Telford, M.J. (2020). Topology-dependent asymmetry in systematic errors affects phylogenetic placement of Ctenophora and Xenacoelomorpha. Science Advances 6, eabc5162. 10.1126/sciadv.abc5162.

30. Redmond, A.K., and McLysaght, A. (2021). Evidence for sponges as sister to all other animals from partitioned phylogenomics with mixture models and recoding. Nat Commun 12, 1783. 10.1038/s41467-021-22074-7.

31. Kapli, P., Natsidis, P., Leite, D.J., Fursman, M., Jeffrie, N., Rahman, I.A., Philippe, H., Copley, R.R., and Telford, M.J. (2021). Lack of support for Deuterostomia prompts reinterpretation of the first Bilateria. Science Advances 7, eabe2741. 10.1126/sciadv.abe2741.

32. Lartillot, N., and Philippe, H. (2004). A Bayesian Mixture Model for Across-Site Heterogeneities in the Amino-Acid Replacement Process. Molecular Biology and Evolution 21, 1095–1109. 10.1093/molbev/msh112.

33. Lartillot, N., Brinkmann, H., and Philippe, H. (2007). Suppression of long-branch attraction artefacts in the animal phylogeny using a site-heterogeneous model. BMC Evolutionary Biology 7, S4. 10.1186/1471-2148-7-S1-S4.

34. Dayhoff, M. O., Schwartz, R. M., and Orcutt, B.C. (1978). A model of evolutionary change in proteins. In Atlas of Protein Sequence and Structure (National Biomedical Research Foundation, Washington DC), pp. 345–352.

35. Foster, P.G., Schrempf, D., Szöllősi, G.J., Williams, T.A., Cox, C.J., and Embley, T.M. (2022). Recoding Amino Acids to a Reduced Alphabet may Increase or Decrease Phylogenetic Accuracy. Systematic Biology, syac042. 10.1093/sysbio/syac042.

36. Giacomelli, M., Rossi, M.E., Lozano-Fernandez, J., Feuda, R., and Pisani, D. (2022). Resolving tricky nodes in the tree of life through amino acid recoding. iScience 25, 105594. 10.1016/j.isci.2022.105594.

37. Feuda, R., Dohrmann, M., Pett, W., Philippe, H., Rota-Stabelli, O., Lartillot, N., Wörheide, G., and Pisani, D. (2017). Improved Modeling of Compositional Heterogeneity Supports Sponges as Sister to All Other Animals. Current Biology 27, 3864–3870.e4. 10.1016/j.cub.2017.11.008.

38. Susko, E., and Roger, A.J. (2007). On Reduced Amino Acid Alphabets for Phylogenetic Inference. Molecular Biology and Evolution 24, 2139–2150. 10.1093/molbev/msm144.

39. Hernandez, A.M., and Ryan, J.F. (2021). Six-State Amino Acid Recoding is not an Effective Strategy to Offset Compositional Heterogeneity and Saturation in Phylogenetic Analyses. Systematic Biology 70, 1200–1212. 10.1093/sysbio/syab027.

40. Kosiol, C., Goldman, N., and H. Buttimore, N. (2004). A new criterion and method for amino acid classification. Journal of Theoretical Biology 228, 97–106. 10.1016/j.jtbi.2003.12.010.

41. Siu-Ting, K., Torres-Sánchez, M., San Mauro, D., Wilcockson, D., Wilkinson, M., Pisani, D., O’Connell, M.J., and Creevey, C.J. (2019). Inadvertent Paralog Inclusion Drives Artifactual Topologies and Timetree Estimates in Phylogenomics. Mol Biol Evol 36, 1344–1356. 10.1093/molbev/msz067.

42. Philippe, H., Vienne, D.M. de, Ranwez, V., Roure, B., Baurain, D., and Delsuc, F. (2017). Pitfalls in supermatrix phylogenomics. European Journal of Taxonomy 283, 1–25. 10.5852/ejt.2017.283.

43. 43. Schiffer, P.H., Natsidis, P., Leite, D.J., Robertson, H., Lapraz, F., Marlétaz, F., Fromm, B., Baudry, L., Simpson, F., Høye, E., et al. (2022). The slow evolving genome of the xenacoelomorph worm Xenoturbella bocki. 2022.06.24.497508. 10.1101/2022.06.24.497508.

44. Gehrke, A.R., Neverett, E., Luo, Y.-J., Brandt, A., Ricci, L., Hulett, R.E., Gompers, A., Ruby, J.G., Rokhsar, D.S., Reddien, P.W., et al. (2019). Acoel genome reveals the regulatory landscape of whole-body regeneration. Science 363, eaau6173. 10.1126/science.aau6173.

45. Martinez, P., Ustyantsev, K., Biryukov, M., Mouton, S., Glasenburg, L., Sprecher, S.G., Bailly, X., and Berezikov, E. (2023). Genome assembly of the acoel flatworm Symsagittifera roscoffensis, a model for research on body plan evolution and photosymbiosis. G3 (Bethesda) 13, jkac336. 10.1093/g3journal/jkac336.

46. Laumer, C.E., Fernández, R., Lemer, S., Combosch, D., Kocot, K.M., Riesgo, A., Andrade, S.C.S., Sterrer, W., Sørensen, M.V., and Giribet, G. (2019). Revisiting metazoan phylogeny with genomic sampling of all phyla. Proceedings of the Royal Society B: Biological Sciences 286, 20190831. 10.1098/rspb.2019.0831.

47. Wilkinson, M., McInerney, J.O., Hirt, R.P., Foster, P.G., and Embley, T.M. (2007). Of clades and clans: terms for phylogenetic relationships in unrooted trees. Trends in Ecology & Evolution 22, 114–115. 10.1016/j.tree.2007.01.002.

48. Shimodaira, H. (2002). An Approximately Unbiased Test of Phylogenetic Tree Selection. Systematic Biology 51, 492–508. 10.1080/10635150290069913.

49. Pisani, D., Pett, W., Dohrmann, M., Feuda, R., Rota-Stabelli, O., Philippe, H., Lartillot, N., and Wörheide, G. (2015). Genomic data do not support comb jellies as the sister group to all other animals. Proceedings of the National Academy of Sciences 112, 15402–15407. 10.1073/pnas.1518127112.

50. DeSalle, R., Narechania, A., and Tessler, M. (2023). Multiple outgroups can cause random rooting in phylogenomics. Molecular Phylogenetics and Evolution 184, 107806. 10.1016/j.ympev.2023.107806.

51. Criscuolo, A., and Gribaldo, S. (2010). BMGE (Block Mapping and Gathering with Entropy): a new software for selection of phylogenetic informative regions from multiple sequence alignments. BMC Evolutionary Biology 10, 210. 10.1186/1471-2148-10-210.

52. Lartillot, N., Lepage, T., and Blanquart, S. (2009). PhyloBayes 3: a Bayesian software package for phylogenetic reconstruction and molecular dating. Bioinformatics 25, 2286– 2288. 10.1093/bioinformatics/btp368.

53. Szánthó, L.L., Lartillot, N., Szöllősi, G.J., and Schrempf, D. (2023). Compositionally Constrained Sites Drive Long-Branch Attraction. Systematic Biology, syad013. 10.1093/sysbio/syad013.

54. Yang, Z. (1997). How often do wrong models produce better phylogenies? Molecular Biology and Evolution 14, 105–108. 10.1093/oxfordjournals.molbev.a025695.

55. Simion, P., Philippe, H., Baurain, D., Jager, M., Richter, D.J., Franco, A.D., Roure, B., Satoh, N., Quéinnec, É., Ereskovsky, A., et al. (2017). A Large and Consistent Phylogenomic Dataset Supports Sponges as the Sister Group to All Other Animals. Current Biology 27, 958–967. 10.1016/j.cub.2017.02.031.

56. Redmond, A.K., and McLysaght, A. (2023). Reply to: Available data do not rule out Ctenophora as the sister group to all other Metazoa. Nat Commun 14, 710. 10.1038/s41467-023-36152-5.

57. Telford, M.J. (2008). Xenoturbellida: The fourth deuterostome phylum and the diet of worms. genesis 46, 580–586. 10.1002/dvg.20414.

58. Jondelius, U., Ruiz-Trillo, I., Baguñà, J., and Riutort, M. (2002). The Nemertodermatida are basal bilaterians and not members of the Platyhelminthes. Zoologica Scripta 31, 201–215. 10.1046/j.1463-6409.2002.00090.x.

59. Ruiz-Trillo, I., Paps, J., Loukota, M., Ribera, C., Jondelius, U., Baguñà, J., and Riutort, M. (2002). A phylogenetic analysis of myosin heavy chain type II sequences corroborates that Acoela and Nemertodermatida are basal bilaterians. Proceedings of the National Academy of Sciences 99, 11246–11251. 10.1073/pnas.172390199.

60. Ruiz-Trillo, I., Riutort, M., Littlewood, D.T.J., Herniou, E.A., and Baguñà, J. (1999). Acoel Flatworms: Earliest Extant Bilaterian Metazoans, Not Members of Platyhelminthes. Science 283, 1919–1923. 10.1126/science.283.5409.1919.

61. Mongiardino Koch, N. (2021). Phylogenomic Subsampling and the Search for Phylogenetically Reliable Loci. Molecular Biology and Evolution 38, 4025–4038. 10.1093/molbev/msab151.

62. Fernández, R., Gabaldon, T., and Dessimoz, C. (2020). Orthology: Definitions, Prediction, and Impact on Species Phylogeny Inference. In Phylogenetics in the Genomic Era (No commercial publisher | Authors open access book), p. pp.2.4:1--2.4:14.

63. Ranwez, V., and Chantret, N.N. (2020). Strengths and Limits of Multiple Sequence Alignment and Filtering Methods. In Phylogenetics in the Genomic Era (No commercial publisher | Authors open access book,), p. p.2.2:1–2.2:36.

64. Tan, G., Muffato, M., Ledergerber, C., Herrero, J., Goldman, N., Gil, M., and Dessimoz, C. (2015). Current Methods for Automated Filtering of Multiple Sequence Alignments Frequently Worsen Single-Gene Phylogenetic Inference. Systematic Biology 64, 778– 791. 10.1093/sysbio/syv033.

65. Francis, W.R., and Canfield, D.E. (2020). Very few sites can reshape the inferred phylogenetic tree. PeerJ 8, e8865. 10.7717/peerj.8865.

66. Li, Y., Shen, X.-X., Evans, B., Dunn, C.W., and Rokas, A. (2021). Rooting the Animal Tree of Life. Molecular Biology and Evolution 38, 4322–4333. 10.1093/molbev/msab170.

67. Philippe, H., Brinkmann, H., Lavrov, D.V., Littlewood, D.T.J., Manuel, M., Wörheide, G., and Baurain, D. (2011). Resolving Difficult Phylogenetic Questions: Why More Sequences Are Not Enough. PLOS Biology 9, e1000602. 10.1371/journal.pbio.1000602.

68. Simion, P., Delsuc, F., and Philippe, H. (2020). To What Extent Current Limits of Phylogenomics Can Be Overcome? In Phylogenetics in the Genomic Era (No commercial publisher | Authors open access book), p. pp.2.1:1--2.1:34.

69. Lozano-Fernandez, J. (2022). A Practical Guide to Design and Assess a Phylogenomic Study. Genome Biology and Evolution 14, evac129. 10.1093/gbe/evac129.

70. Fleming, J.F., Valero-Gracia, A., and Struck, T.H. (2023). Identifying and addressing methodological incongruence in phylogenomics: A review. Evolutionary Applications 16, 1087–1104. 10.1111/eva.13565.

71. Kumar, S. (2022). Embracing Green Computing in Molecular Phylogenetics. Molecular Biology and Evolution 39, msac043. 10.1093/molbev/msac043.

72. Telford, M.J., and Copley, R.R. (2011). Improving animal phylogenies with genomic data. Trends in Genetics 27, 186–195. 10.1016/j.tig.2011.02.003.

73. Rokas, A., and Holland, P.W.H. (2000). Rare genomic changes as a tool for phylogenetics. Trends in Ecology & Evolution 15, 454–459. 10.1016/S0169-5347(00)01967-4.

74. Schultz, D.T., Haddock, S.H.D., Bredeson, J.V., Green, R.E., Simakov, O., and Rokhsar, D.S. (2023). Ancient gene linkages support ctenophores as sister to other animals. Nature, 1–8. 10.1038/s41586-023-05936-6.

75. Achatz, J.G., Chiodin, M., Salvenmoser, W., Tyler, S., and Martinez, P. (2013). The Acoela: on their kind and kinships, especially with nemertodermatids and xenoturbellids (Bilateria incertae sedis). Org Divers Evol 13, 267–286. 10.1007/s13127-012-0112-4.

76. Nguyen, L.-T., Schmidt, H.A., von Haeseler, A., and Minh, B.Q. (2015). IQ-TREE: a fast and effective stochastic algorithm for estimating maximum-likelihood phylogenies. Mol Biol Evol 32, 268–274. 10.1093/molbev/msu300.

77. Minh, B.Q., Nguyen, M.A.T., and von Haeseler, A. (2013). Ultrafast approximation for phylogenetic bootstrap. Mol Biol Evol 30, 1188–1195. 10.1093/molbev/mst024.

78. Kalyaanamoorthy, S., Minh, B.Q., Wong, T.K.F., von Haeseler, A., and Jermiin, L.S. (2017). ModelFinder: fast model selection for accurate phylogenetic estimates. Nat Methods 14, 587–589. 10.1038/nmeth.4285.

79. Kishino, H., Miyata, T., and Hasegawa, M. (1990). Maximum likelihood inference of protein phylogeny and the origin of chloroplasts. J Mol Evol 31, 151–160. 10.1007/BF02109483.

80. Le, S.Q., and Gascuel, O. (2008). An Improved General Amino Acid Replacement Matrix. Molecular Biology and Evolution 25, 1307–1320. 10.1093/molbev/msn067.

81. Yang, Z. (1994). Maximum likelihood phylogenetic estimation from DNA sequences with variable rates over sites: Approximate methods. J Mol Evol 39, 306–314. 10.1007/BF00160154.

82. Si Quang, L., Gascuel, O., and Lartillot, N. (2008). Empirical profile mixture models for phylogenetic reconstruction. Bioinformatics 24, 2317–2323. 10.1093/bioinformatics/btn445.

83. Wang, H.-C., Minh, B.Q., Susko, E., and Roger, A.J. (2018). Modeling Site Heterogeneity with Posterior Mean Site Frequency Profiles Accelerates Accurate Phylogenomic Estimation. Systematic Biology 67, 216–235. 10.1093/sysbio/syx068.

84. Ly-Trong, N., Naser-Khdour, S., Lanfear, R., and Minh, B.Q. (2022). AliSim: A Fast and Versatile Phylogenetic Sequence Simulator for the Genomic Era. Molecular Biology and Evolution 39, msac092. 10.1093/molbev/msac092.

85. Minh, B.Q., Schmidt, H.A., Chernomor, O., Schrempf, D., Woodhams, M.D., von Haeseler, A., and Lanfear, R. (2020). IQ-TREE 2: New Models and Efficient Methods for Phylogenetic Inference in the Genomic Era. Mol Biol Evol 37, 1530–1534. 10.1093/molbev/msaa015.

86. Huerta-Cepas, J., Serra, F., and Bork, P. (2016). ETE 3: Reconstruction, Analysis, and Visualization of Phylogenomic Data. Molecular Biology and Evolution 33, 1635–1638. 10.1093/molbev/msw046.

