## Supplementary Figures for "Acoelomorph flatworm monophyly is a severe long branch-attraction artefact obscuring a clade of Acoela and Xenoturbellida"

**Anthony K Redmond**
Smurfit Institute of Genetics, Trinity College Dublin, Dublin 2, Ireland

### Supplementary Figures:

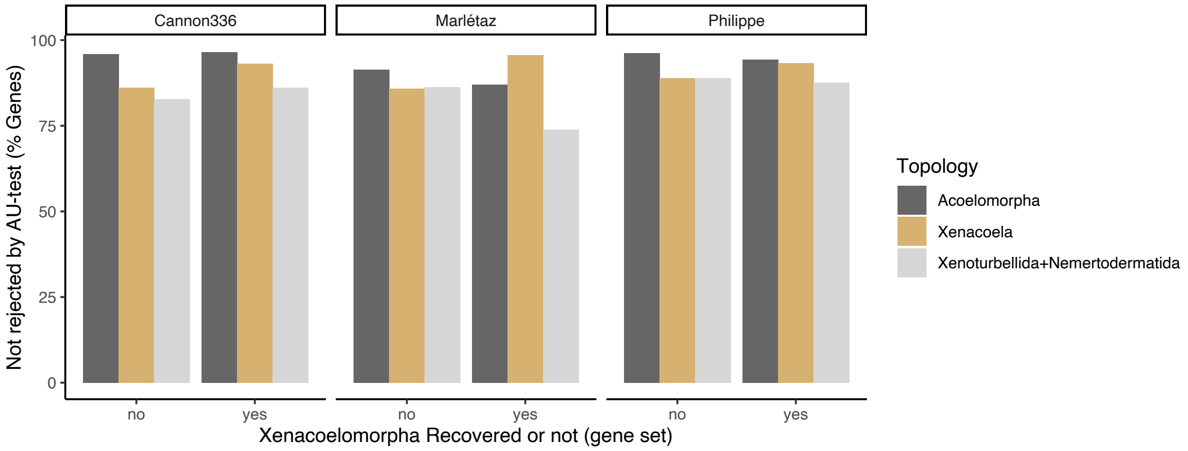


**Figure S1. Proportion of genes that either recover (yes; the 'X' main dataset genes) or do not recover (no) Xenacoelomorpha which do not reject any of three possible xenacoelomorph relationships.**

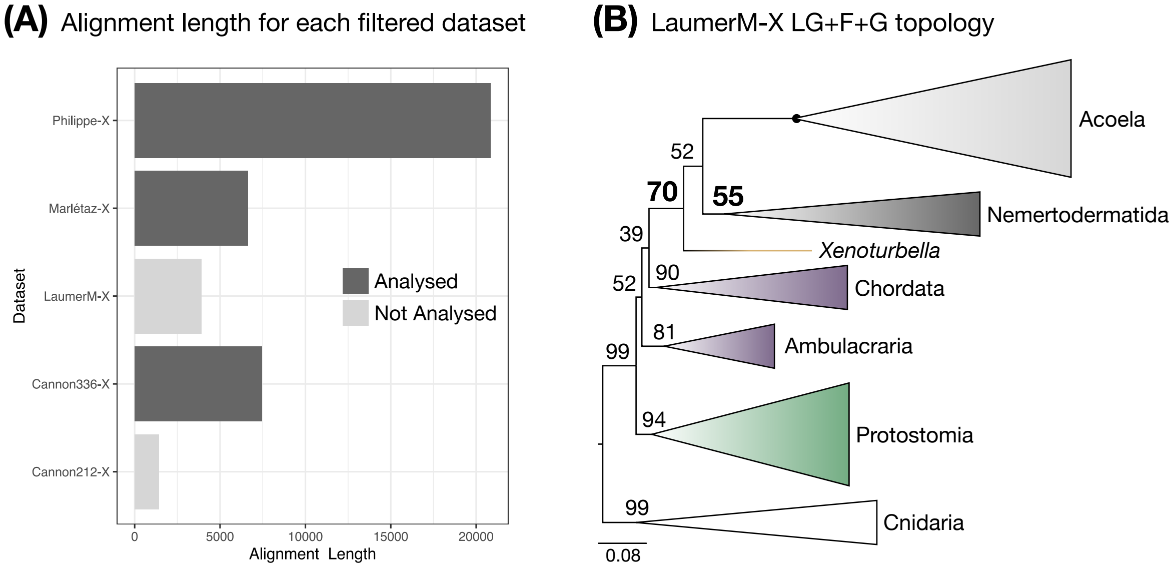


**Figure S2. Further details on dataset exclusion/inclusion.** (**A**) Alignment lengths for the three main datasets as well as for the filtered version of the Laumer pan-metazoan matrix and the main dataset from Cannon et al (2019) study (LaumerM-X and Cannon212-X) that were not employed in further analyses. **(B)** IQ-TREE LG+F+G maximum likelihood consensus tree for the LaumerM-X dataset. Note that low support values for key nodes (Xenacoelomorpha=70% and Nemertodermatida=55% UFBoot) adjacent to Acoelomorpha/Xenacoela (which itself is near equivocal) are shown in bold. Major lineages are collapsed. Black circles indicate nodes with maximal support.

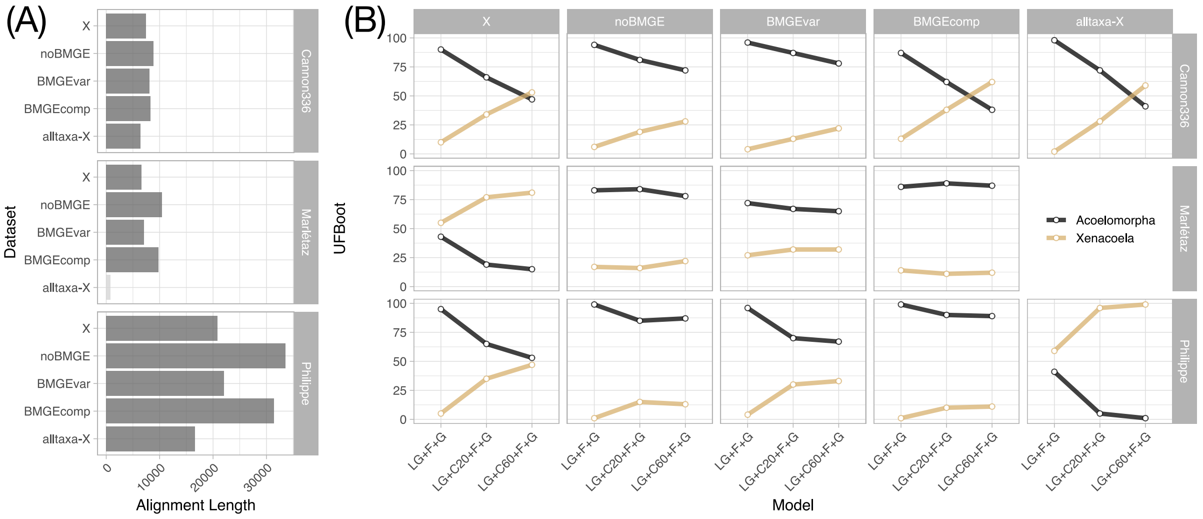


**Figure S3. Analyses of datasets without taxon subsampling or without site-stripping of compositionally biased and/or unexpectedly variable sites.** (**A**) Alignment lengths for datasets with various filtering levels/approaches. The lighter coloured Marlétaz-alltaxa-X dataset was not consider for further analysis due to its short length. (**B)** UFBoot support value trends under the site homogeneous LG+F+G and the precomputed site-heterogeneous LG+C20+F+G and LG+C60+F+G models for Acoelomorpha and Xenacoela compared for datasets with various filtering levels/approaches. “X” = the main ‘-X’ datasets with subsampled taxa, and BMGE trimming of both compositionally biased and unexpectedly variable sites used in this study; “noBMGE” = subsampled taxa dataset but without any BMGE site-stripping (compositionally biased and unexpectedly variable sites are both still included); “BMGEvar” = subsampled taxa dataset with BMGE site-stripping of unexpectedly variable sites only; “BMGEcomp” = subsampled taxa dataset with BMGE site-stripping of compositionally biased sites only; “alltaxa-X” = dataset without taxon subsampling but with full BMGE site-stripping of both compositionally biased and unexpectedly variable sites (this leads to fewer retained sites than the main ‘-X’ datasets, likely due to increased propensity for compositional bias and high variability from inclusion of additional taxa).

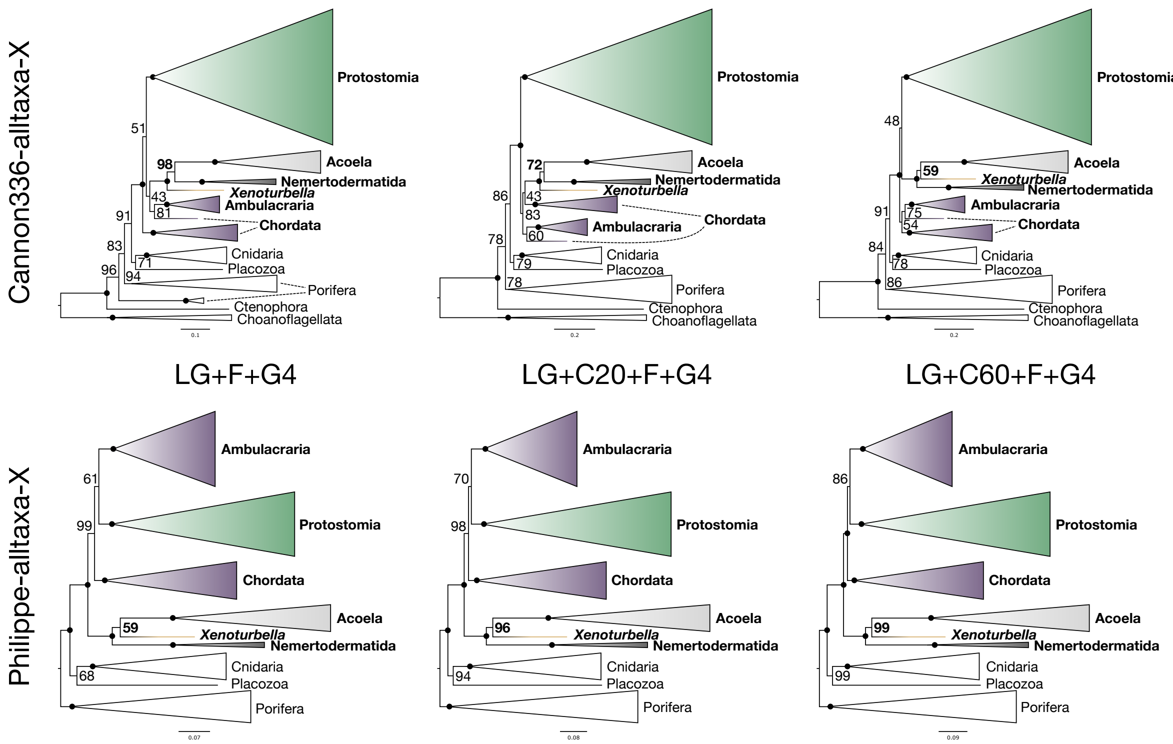


**Figure S4. IQ-TREE maximum likelihood consensus trees under the site homogeneous LG+F+G and the precomputed site-heterogeneous LG+C20+F+G and LG+C60+F+G models for the filtered Cannon336 and Philippe datasets without taxon subsampling.** Major lineages are collapsed and UFBoot support values are shown for all remaining visible branches. Support for Acoelomorpha/Xenacoela is shown in bold. Black circles indicate nodes with maximal support.
